# Geographic variation in hemocyte diversity and phagocytic propensity shows a diffuse genomic signature in the green veined white butterfly

**DOI:** 10.1101/790782

**Authors:** Naomi L.P. Keehnen, Lisa Fors, Peter Järver, Anna-Lena Spetz, Sören Nylin, Ulrich Theopold, Christopher W. Wheat

## Abstract

Insects rely on their innate immune system to successfully mediate complex interactions with their internal microbiota, as well as the microbes present in the environment. Given the variation in microbes across habitats, the challenges to respond to them is likely to result in local adaptation in the immune system. Here we focus upon phagocytosis, a mechanism by which pathogens and foreign particles are engulfed in order to be contained, killed and processed for antigen presentation. We investigated the phenotypic and genetic variation related to phagocytosis, in two allopatric populations of the butterfly *Pieris napi*. We found that the populations differ in their hemocyte composition, and overall phagocytic capability, driven by the increased phagocytic propensity of each cell type. However, no evidence for divergence in phagocytosis-related genes was observed, though an enrichment of genes involved in glutamine metabolism was found, which have recently been linked to immune cell differentiation in mammals.

## Introduction

Comparisons between populations has long been a powerful tool used by evolutionary biologist to identify both the action and potential targets of natural selection in wild populations (e.g. Endler 1977). While common garden experiments can remove environmental effects, deeper dissection of phenotypic differences remains challenging. Consider physiological differences, where phenotype differences might arise from changes in the amount, size or performance of cells or organs. Disentangling these relative contributions to phenotypic differences is important, as the targets of selection are expected to be very different when the target is the genes directly involved in the phenotype vs. cell proliferation or differentiation. Here we draw attention to this issue in the field of ecological immunity, as immune performance assays in natural populations often only focus upon phenotypic differences in immune performance without distinguishing between these two different causal routes. However, such insights are needed if we are to understand the microevolutionary dynamics of ecological immunity, as it informs upon whether the immune pathways or the proliferation of immune cells are the target of selection on immune performance.

Insects are ubiquitous, occurring in a wide variety of environments. As such they are exposed to a wide variety of parasites and microbes, both from their environment, as well as within themselves, i.e. their microbiome. These interactions with pathogens and parasites are mediated by their immune system. Natural variation in immunity is widespread among individuals (Kurtz et al. 2000), populations (Kraaijeveld and Van Alphen 1995; Tinsley et al. 2006; Corby-Harris & Promislow, 2008), and species (Palmer et al. 2018). This variation could be a response to geographic variation in biotic characteristics, e.g. bacterial density or parasite prevalence in the environment or within themselves (Corby-Harris & Promislow, 2008; Fors et al. 2016; Coon et al. 2016; Chaplinska et al., 2016), or abiotic factors such as temperature and nutrition (Ferguson & Read 2002; Blanford et al. 2003; Mitchell et al. 2005; Lazzaro et al. 2008). These variable selection pressures can potentially shape and drive adaptations in the immune system (Schmid-Hempel, 2003; Vogelweith et al. 2013; Buchon et al. 2009; Kraaijeveld & Godfray 2008; Khan et al. 2017).

The insect immune system can be divided into humoral and cellular defence responses. Humoral responses involve the production of antimicrobial peptides and enzymes of the phenoloxidase cascade (Charroux and Royet, 2010). The cellular responses, often the largest proportion of the immune response (Haine et al. 2008), are mediated by the blood cells (hemocytes), and include encapsulation, phagocytosis, and nodulation. Phagocytosis is one of the most widely conserved defence mechanisms against microorganisms. The process begins when pathogens or dead cells are recognized by pattern recognition receptors (PRRs), which bind the particle and activate intracellular cascades leading to the pathogen being engulfed into the phagosome, after which it is eliminated (Strand, 2008; Stuart & Ezekowitz 2008).

To our knowledge, studies on variation in phagocytic capabilities of invertebrates are scarce, but variation does exist at several evolutionary scales. At the macroevolutionary scale, comparative analysis across six arthropods found variation in phagocytic performance that was correlated with body size, as organisms with fewer hemocytes were smaller, yet had higher phagocytic capabilities (Oliver et al. 2011). Studies of closely related species have also been found to differ in their phagocytic activity, as two closely related heliothine moth species, *Heliothis irescens* and *H. subflexa*, differ in their phagocytic activity hypothesized to be a result of their hemocyte composition differences (Barthel et al. 2014). In addition, phagocytic performance differences have been observed among different castes of the eusocial honey bee (*Apis mellifera*) (Hystad et al. 2017). Lastly, differences between sexes have also been observed, with females found better at phagocytosing than males (Kurtz et al. 2000). However, we have been unable to find studies investigating population level differences in phagocytosis.

While the literature at the phenotypic level is sparse, genome signatures of selection have identified genes having functional annotations related to phagocytosis. Phagocytosis-associated genes, across a wide variety of insects, show fast rates of evolution and high population differentiation (Sackton et al. 2007; Waterhouse et al. 2007; Crawford et al. 2010, Chavez-Galarza et al. 2013; Erler et al. 2014). In *Drosophila*, genes encoding PRRs specific to phagocytosis appear to evolve more rapidly and display high levels of population differentiation (Sackton et al. 2007; Juneja & Lazarro 2010; Early et al. 2017). However, whether it was selection upon the phagocytic performance of these genes that drove this increased rate of evolutionary change remains to be investigated.

In sum, while there is evidence for both phenotypic and genotypic variation in the wild in relation to phagocytosis, little is known about how this genetic variation interacts with variation on the phenotypic level, and whether this affects phagocytic capabilities. Furthermore, to our knowledge, the majority of the studies looking at phagocytosis has been performed at the species level, reflecting a lack of information obtained from spatial variability of immune defences within species and between populations in the wild.

Here we work to integrate these top down (phenotype level) and bottom up (genomic patterns) perspectives by investigating the phenotypic and genetic variation related to phagocytosis in populations of a Lepidopteran butterfly, the Green veined White (*Pieris napi*). We aim to characterize differences between the populations in their innate immune responses, and then assess whether this correlates with divergence in relevant immune genes between these populations. Specifically, we investigate phagocytic performance variation at both the phenotypic and genomic level between two populations of the Green veined White butterfly (*Pieris napi*), one from Aiguamolls in northern Spain, the other from Abisko in northern Sweden. *P. napi* is a common and widespread butterfly that feeds exclusively upon Brassicales plants, with limited dispersal and therefore low gene flow among populations (Porter and Geiger, 1995). This species produces between one (northern Europe) to four generations (southern Europe) per year depending on local conditions, reflecting local adaptation along latitude (Pruisscher et al. 2017). This variation in life history makes it an interesting species to investigate potential local adaptation of the immune system. Using common garden reared individuals and several different measures of phagocytic performance, we find significant differences in the phagocytic performance between the two populations, as well as between sexes. Furthermore, we characterized the phagocyting cell types, and their relative performance, finding that both cell composition and propensity to phagocytose differ between the two populations. Finally, using a population genomics analysis, we investigated to what extent these populations differ in genes of the recognized phagocytic pathway, finding that there is little divergence or turnover at these genes. Instead, genes involved in proteolysis, metabolic and catabolic processes displayed differences providing candidates for further studies.

## Material and Methods

### Study species

*P. napi* adult females were caught in spring 2016 from Spain (Parc dels Aiguamolls de l’Empordà, north-east of Barcelona, 42.23°N, 3.10°E) and northern Sweden (Abisko 68.36°N, 18.79°E and Kiruna, 67.87°N, 20.17°E). Butterflies were kept separately in 1-L cups to lay eggs on *Alliaria petiolata*. Plants were replaced daily. Offspring were reared under 8:16 hours Light:Dark, and 17°C to ensure diapause development in both populations (Pruisscher et al. 2017), and placed into 2°C 0:24 L:D for overwintering. They were kept per 4 in 1-L cups, and fed a mixture of wild *A. petiolata* and *Armoracia rusticana*. In spring 2017 the diapausing pupae were taken out of the cold, and reared for another generation as described above, only crossing between unrelated families. Larvae were observed during molting stages, and once they reached the final, 5^th^ instar they were weighed and assayed.

### Total hemocyte count

For the total hemocyte count (THC) assay 20 Spanish and 18 Swedish 5^th^ instar larvae (F:M) were used. All larvae were sampled on the first day of the 5^th^ instar, of approximately similar weight (weighed at a maximum of 2 hours prior to the experiment). The larvae were bled by cutting the second proleg and 8 µl hemolymph directly collected and added to 152 µl anticoagulant buffer (dilution 1:20), prepared according to Firlej et al. 2012 (62mM NaCl, 100 mM glucose, 10 mM EDTA, 30 mM trisodium citrate and 26 mM citric acid). Samples were gently mixed and 10 µl directly transferred to each side of a hemocytometer (Neubauer chamber, 0.1 mm depth). Hemocytes were counted in 10 squares for each individual and the cell concentration in 1 µl of hemolymph calculated by dividing (the number of cells x10) by (the number of squares x dilution).

### Phagocytosis slide assay

To investigate the presence of phagocytic cells, 10 µl suspension of, heat killed *Escherichia coli* (Thermofisher Scientific *E. coli* (K-12 strain) BioParticles™-Alexa Fluor™ 488 conjugate, a green-fluorescent dye) was injected into early 5^th^ instar larvae (22 larva per population, 11 of each sex). The bacteria were injected on the ventral side behind the second proleg, using a glass capillary needle. After injection, the larvae were kept in separate containers with *Alliaria petiolate* ad libitum for 4 hours (at room temperature RT) before sample preparation. After 4 hours of incubation, the larvae were bled by cutting the second proleg, whereupon 5 µl hemolymph/ individual was collected and added to separate vials containing 300 µl Phosphate buffered saline (PBS) mixed with a small amount of phenylthiourea (PTU, Sigma Aldrich) and with 5% Newborn Calf Serum (NBCS, Biowest). Individuals showing signs of open wounds were excluded to avoid the risk of unequal bacteria concentration. Two hemocyte samples were prepared from each individual larva. For each sample, a 30 µl drop of the mixture (hemolymph + PBS) was placed on a multi-spot microscope slide (SM-011, Hendley, Loughton, U.K.) and left to adhere for 20 minutes in a humid chamber at room temperature (RT). After adhesion excessive hemolymph was gently removed with a pipette and the remaining cell monolayer was fixed with 4% PFA + PBS for 10 minutes. After fixation cells were washed three times with PBS for 5 minutes. To distinguish morphological features and reveal the nuclei the cells were treated with Phalloidin Rhodamine (Biotium, 3:1000 dilution in PBS) together with blue-fluorescent nucleic acid stain DAPI (Sigma-Aldrich, 1:1000 dilution in PBS) for 10 minutes in darkness, RT. After DNA-staining the cells were washed three times with PBS for 5 minutes and mounted in Fluoromount-G (SouthernBiotech) mounting media. All samples were studied in a Zeiss Axioplan2, phase contrast, epifluorescent microscope connected to a Hamamatsu camera with Axio Vision 4.6. Eight random images per individual were taken and used for differential hemocyte counts and assessment of phagocytic capacity. Additionally, most samples were also studied in a Zeiss LSM 780 confocal microscope for acquisition of Z-stack images to further verify bacterial uptake.

### Quenching with trypan blue

To verify that the Alexa Fluor 488, heat killed *E. coli* was ingested by the hemocytes and not just attached to the surface of cells, additional tests were performed using trypan blue to quench the signal from extracellular bacteria. Early 5^th^ instar larvae were bled and 10 µl hemolymph/ individual was collected and added to vials with 300 µl PBS mixed with 5% NBCS and a small amount of PTU as above. 2,5 µl suspension of *E. coli* was added to each vial and the solution mixed gently but thoroughly, whereupon 30 µl of the mixture was placed on a multi-spot microscope slide and left to adhere for 20 minutes in a humid chamber at room temperature. Similar samples were prepared from injected individuals (without the addition of *E. coli* after bleeding). Just before the samples were studied in the microscope, 20 µl of each sample was removed and 10 µl freshly prepared 0, 4% trypan blue was added. After 1-2 minutes incubation, 10-15 µl of the mixture was removed with a pipette and a coverslip gently placed on top of the samples.

### Phagocytosis Flow cytometry assay

To assess phagocytic capacity, *in vitro* samples of hemocytes treated with pHrodo conjugated *E. coli* (pHrodo™ Green E. coli BioParticles® Conjugate, Thermo Fisher Scientific) were prepared. The pHrodo Green conjugates are non-fluorescent outside the cell but fluoresce brightly green as pH decreases from neutral to acidic in the phagosomes, thereby providing a robust quantitative measure of phagocytic activity. A total of 74 Spanish and 48 Swedish larvae were used. All larvae used were in early 5^th^ instar and of approximately the same size (weighed at a maximum of 2 hours prior to the experiment). The larvae were bled by cutting the second proleg, whereupon 25 µl hemolymph/ individual was collected and added to separate vials containing 300 µl Phosphate buffered saline (PBS) with 5% Newborn Calf Serum (NBCS, Biowest). The samples were then gently mixed with 50 µl pHrodo conjugated *E. coli* and incubated for 3 hours (darkness, RT). Two control samples were prepared for each trial (one male and one female), treated with the same amount of *E. coli* but incubated on ice to inhibit the phagocytic process. After incubation, each sample was analyzed individually by a Fortessa flow cytometer (BD Biosciences) and the data analysis was performed with FlowJo software (TreeStar).

### Immunostaining of hemocytes with Prophenoloxidase Antiserum

To distinguish the oenocytoids from other cell types an additional test was performed using an antiserum to *Manduca sexta* prophenoloxidase (courtesy of Micheal R Kanost, Department of Biochemistry and Molecular Biophysics Kansas State University). 5^th^ instar larvae were bled by cutting the second proleg, whereupon 10 µl hemolymph/ individual was collected and added to separate vials containing 300 µl Phosphate buffered saline (PBS) with 5% Newborn Calf Serum (NBCS, Biowest). Three hemocyte samples were prepared from each individual larva. For each sample, a 30 µl drop of the mixture (hemolymph + PBS) was placed on a category number 1.5 coverslip (0.17mm) and left to adhere for 1 hour in a humid chamber at room temperature (RT). After adhesion excessive hemolymph was gently removed with a pipette and the remaining cell monolayer was fixed with 4% PFA + PBS for 5 minutes. After fixation cells were washed three times with PBS for 5 minutes, permeabilized with 0,1% Triton X-100 in PBS for 5 minutes and then washed again three times with PBS. Blocking was performed by adding 3% Bovine Serum Albumin (BSA) for 30 min, after which the cells were incubated with the primary antibody (prophenoloxidase antiserum) for 2 hours (darkness, RT). After washing with PBS three times, the cells were incubated with the secondary antibody (Alexa 488, anti-rabbit) for 2 hours (darkness, RT), and then washed as above. To distinguish morphological features and reveal the nuclei the cells were treated with Phalloidin Rhodamine (Biotium, 3:1000 dilution in PBS) together with blue-fluorescent nucleic acid stain DAPI (Sigma-Aldrich, 1:1000 dilution in PBS) for 10 minutes (darkness, RT). After staining the cells were washed four times with PBS for 5 minutes and mounted in Fluoromount-G (SouthernBiotech) mounting media on a regular microscope slide. All samples were studied in a Zeiss LSM 780 confocal microscope.

### Data analysis

All statistical tests were performed using the statistical software R version 3.4.3. In order to correct for the pseudoreplicated nature of the data generalized linear mixed models (lme4 package) were used, including the replicates as random factor. We performed stepwise backwards elimination for each GLM, starting with all interactions, and removing each least-significant term until the AIC and BIC no longer changed. In the analysis of the propensity of a given cell type that was phagocytosing, not all slides contained all types of cells, so for these analyses we performed a GLM using quasibinomial distribution.

### Gene scan and population genetics

Here, we use the chromosomal level genome of *Pieris napi* (Hill et al. 2019). As this genome, as well as other Lepidopteran genomes, have few annotations for phagocytosis genes, we generated annotations using proteins previously indicated to be involved with phagocytosis in *Drosophila melanogaster* (N = 155; Im & Lazzaro 2008) using SPALN v2.1.2 (Gotoh, 2008; Iwata & Gotoh, 2012). To validate the accuracy of the identified phagocytosis genes, we implemented a number of validation steps. First, we used TBLASTN (Altschul, Gish, Miller, Myers, & Lipman, 1990) with the 155 proteins to search the genome assembly, extracting hits with *E*-values < 0.0001 and a bitscore higher than 60. Second, we used MESPA (Neethiraj et al. 2017) to conduct protein searches of the genome for the 155 candidate genes and generate gene models (*n* = 73 were predicted). We then compared the blast hits with the SPALN gene models, to assess gene model accuracy, filter out potential errors and identify potential duplicates. As a last quality control step, each gene model was manually investigated using IGV (Robinson et al., 2011). To be confident in our gene models, average read depth per exon was calculated using coverageBed from Bedtools2 (Quinlan & Hall, 2010) and compared to the coverage of all genes. These approaches resulted in a total of 73 phagocytosis-related genes identified in the genome.

For our population genomics analysis, we used two previously published (Keehnen et al. 2018) Pool-seq datasets originating from the two populations used in this experiment. These two libraries were mapped against the genome using Next-Gen mapper v.04.10. Samtools v1.2 was used to filter the mapped data for mapped paired-end reads, after which a mpileup file was created for further analysis (Li et al., 2009). Popoolation v1.2.2 (Kofler et al. 2011) was used to mask indels using a 5-bp window, centered in the indel (identify-genomic-indel-regions.pl), as well as to mask non-genic regions (create-genewise-sync.pl). Population differentiation was identified using the fixation index (F_ST_), a measure of population differentiation wherein a 0 indicates no differentiation, whereas a value of 1 implies populations have fixed alternative allelic states and are completely differentiated. F_ST_ was calculated within the coding regions of genes, at the single nucleotide polymorphism (SNP) level, as well as estimated among all exons per gene, using Popoolation2 v1.201 (Kofler, Pandey, & Schlötterer, 2011). In addition, F_ST_ was calculated for SNPs located within 5kb up and downstream of genes selected using the slopBed function of bedtools2 (Quinlan & Hall, 2010). To assess whether the F_ST_ patterns of phagocytosis genes differed from the rest of the genome two permutation subset analysis were performed in R (R Core Team, 2016) to control for differences in subset size. First, we estimated the mean F_ST_ of our 73 phagocytosis genes, and then compared this to a permutation test of 10,000 sets of 73 randomly sampled genome-wide genes. Secondly, we estimated the F_ST_ of 10,000 sets of 1906 randomly sampled genome-wide SNPs and compared this distribution of means to the mean of the 1906 phagocytosis SNPs. Finally, a GSEA was performed, this the SNPs with outlier values, i.e. SNPs with a value that that corresponded to the genome-wide 97.5^th^ percentile were used in GO-term enrichment analysis using TopGO v.2.36.0 (Alexa et al. 2006). We also added all of our identified phagocytosis genes to a phagocytic GO term for analysis. In topGO, the nodeSize parameter was set to 5 to remove GO terms which have fewer than five annotated genes, and other parameters were run on default.

## Results

### Total hemocyte count, cell type identification & characterization

In naive animals, the total hemocyte assay revealed that on average, a 5^th^ instar *P. napi* larvae had 14,037 (SD= 7,125) hemocytes. There was no significant difference in total hemocyte count between larvae from different populations or sex (Figure 1A; GLM: P_population_ = 0.43, P_sex_ = 0.45, SM Table 1). However, among animals injected with bacteria for our phagocytosis assay, the total number of hemocytes differed significantly between sexes, with females having a higher number of hemocytes than males (Figure 1B; P = 0.013), but there was no significant difference between the populations (P = 0.092, SM Table 2). Thus, while naïve animals do not differ in total hemocyte counts, differences between the sexes were apparent after immune challenge.

**Figure 1.**
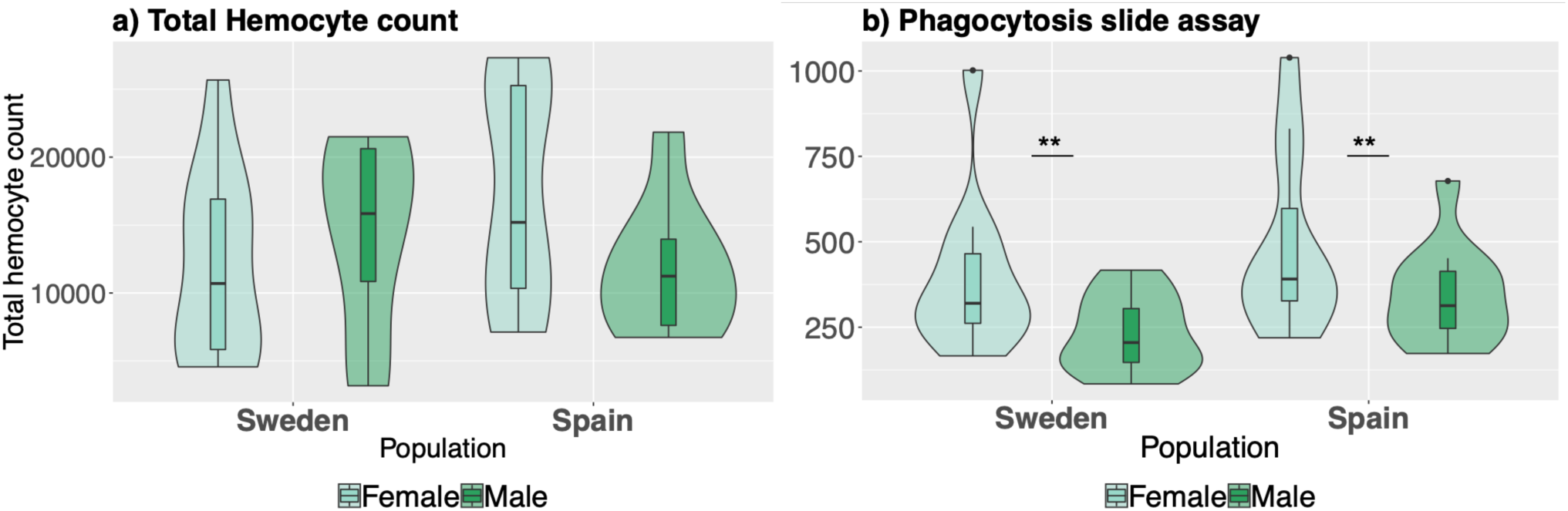
(a) Total hemocyte count of *P. napi* larva per population and sex (N=38 larvae). (b) The total number of hemocytes counted during the phagocytosis slide assay (N=44 larvae).

During our experiments, three cell types were identified to be capable of phagocytosis, and therefore are discussed more in depth. These were granulocytes, plasmatocytes, and (early) oenocytoids (SM Table 3). The majority of the circulating hemocytes were granulocytes. Granulocytes are small, round cells with granular morphology and centralized nucleus. Morphologically they can be identified by their protruding spikes from the regular circle, and they have a weaker phalloidin staining than plasmatocytes (Fig 2A). Plasmatocytes are larger cells having a large elliptic nucleus, with long spikes that protrude in irregular shapes (Fig 2B). They often had weaker nuclear staining but strong phalloidin staining of filaments. Oenocytoids are large, round or slightly elliptic cells with a slightly larger, often not centralized, nucleus than granulocytes and a more homogeneous cytoplasm (Fig. 2C). Furthermore, they appear to have a smoother surface compared to the other cell types. In our studies we observed oenocytoids of very different sizes, representing different stages of the cell type. The phagocytic capacity was mainly detected in the smaller cells, i.e. early or immature oenocytoids (2E), but was not observed in the larger cells, i.e. mature oenocytoids. Oenocytoids identity was further verified by immunostaining with Prophenoloxidase-specific antiserum (2F). The proportion of granulocytes did not differ between the populations (Figure 3a, P = 0.21), or between sexes (P = 0.063, SM Table 4). However, the proportion of plasmatocytes was significantly higher in larvae from Spain (Figure 3b, P = 0.029), but again no effect of sex (P = 0.14, SM Table 5). Finally, there was an interaction between sex and population for the number of oenocytoids (P=0.049, SM Table 6), where males from Sweden had a higher proportion compared to the other sex and population (Figure 3c).

**Figure 2.**
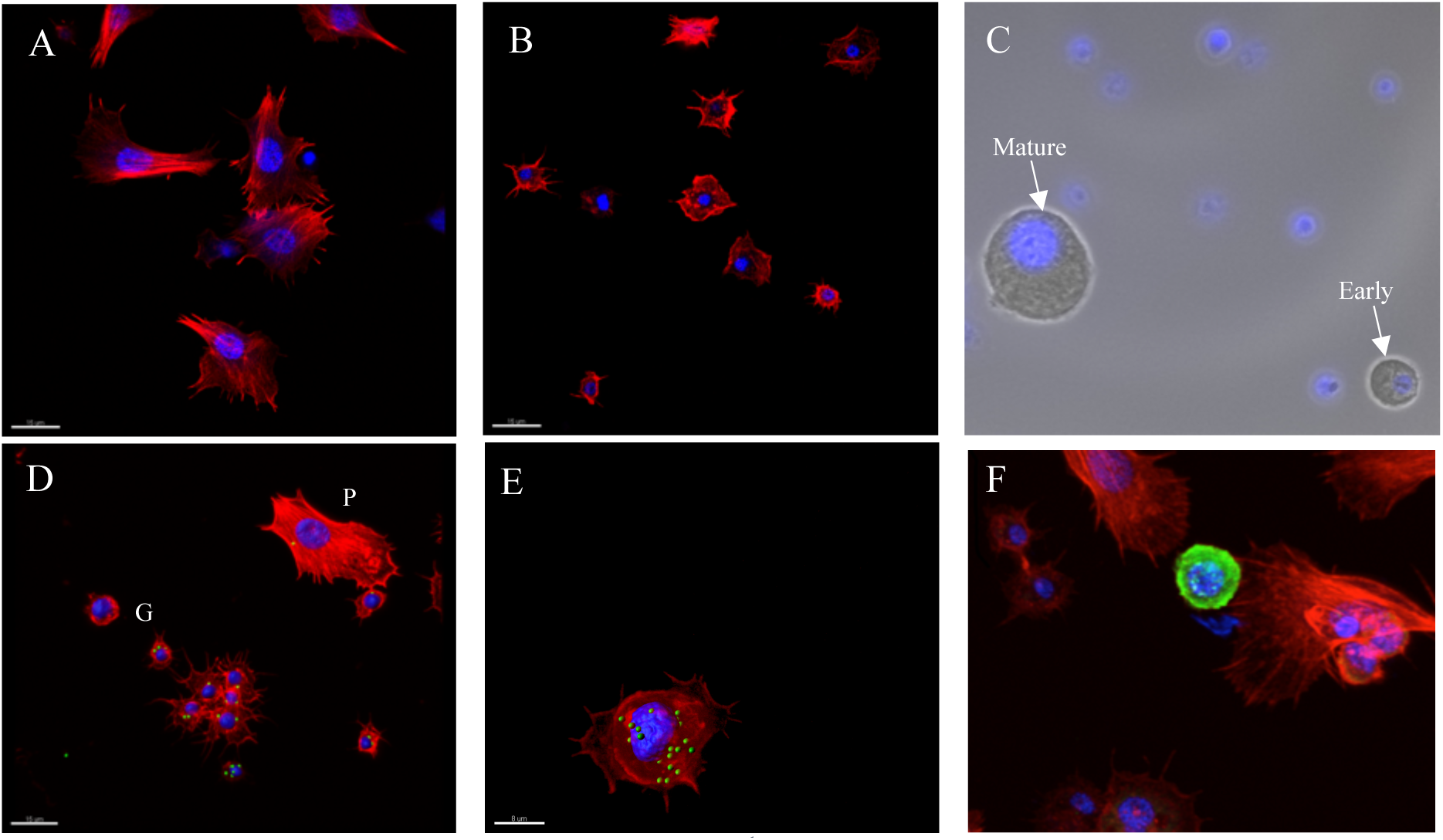
Morphology of the three main types of hemocytes in 5^th^ instar *P. napi* larvae: (A) plasmatocytes, (B) granulocytes, and (C) early and mature oenocytoids. Hemocytes of *P. napi* larvae after in vivo phagocytosis of *E. coli* (AlexaFluor488, green): (D) plasmatocyte (indicated by ^P^) and granulocytes (indicated by ^G^), (E) and early oenocytoid. (F) Staining of an oenocytoid with PPO antibody (green). Cell nuclei are stained with DAPI (blue) and the cytoskeleton of A,B,D,E&F with phalloidin (red). Scalebar = 15 um. Images taken using epifluorescent microscope (C) and confocal microscope (A,B,D,E&F).

**Figure 3.**
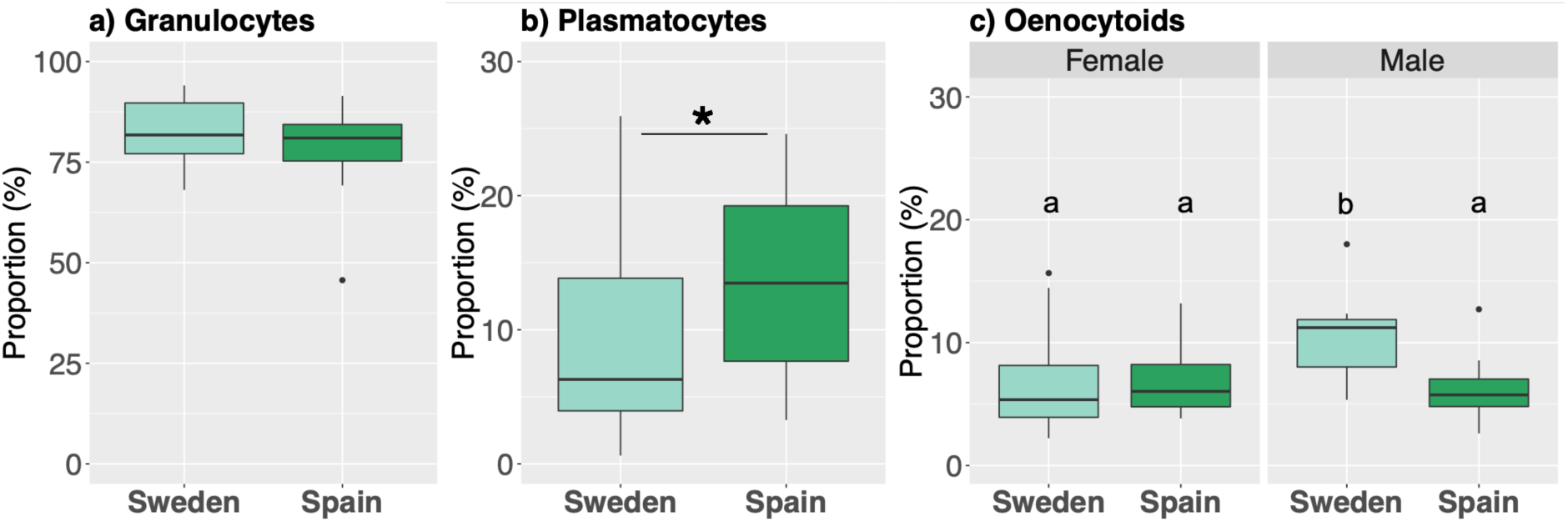
The cell composition of the total number of hemocytes during the phagocytosis assay as measured via imaging (N=44 larvae). The most abundant cell types are granulocytes, followed by plasmatocytes and oenocytoids. The whiskers represent the largest and smallest values within 1.5* interquartile range. The letters in plot c denote significance relationship.

### Phagocyte differences between populations

Next, we investigated the number of phagocytes (i.e. cells that engulfed bacteria) between the populations, using our phagocytic assay (slide assay) and found that they differed significantly (P = 0.0003, SM Table 7). Swedish larvae had significantly higher proportion of phagocytes (80.45%) compared to the Spanish (66.90%). Furthermore, males on average had a higher proportion of phagocytes compared to females (Figure 4a, P= 0.029, SM Table 7). Using an independent phagocytic assay (flow cytometry; gating strategy and examples of staining are shown in Figure S2), a similar population pattern was found, i.e. the Swedish had a significantly higher proportion of phagocytes compared to the Spanish (P < 0.001, SM Table 8). However, the sex differences were not observed using this method (Figure 4b, P = 0.89, SM Table 8). Thus, we confirmed significant differences between the two populations in the phagocytic capacity of challenged individuals using two independent methods. However, the sex difference was only detected using the phagocytic slide assay.

**Figure 4.**
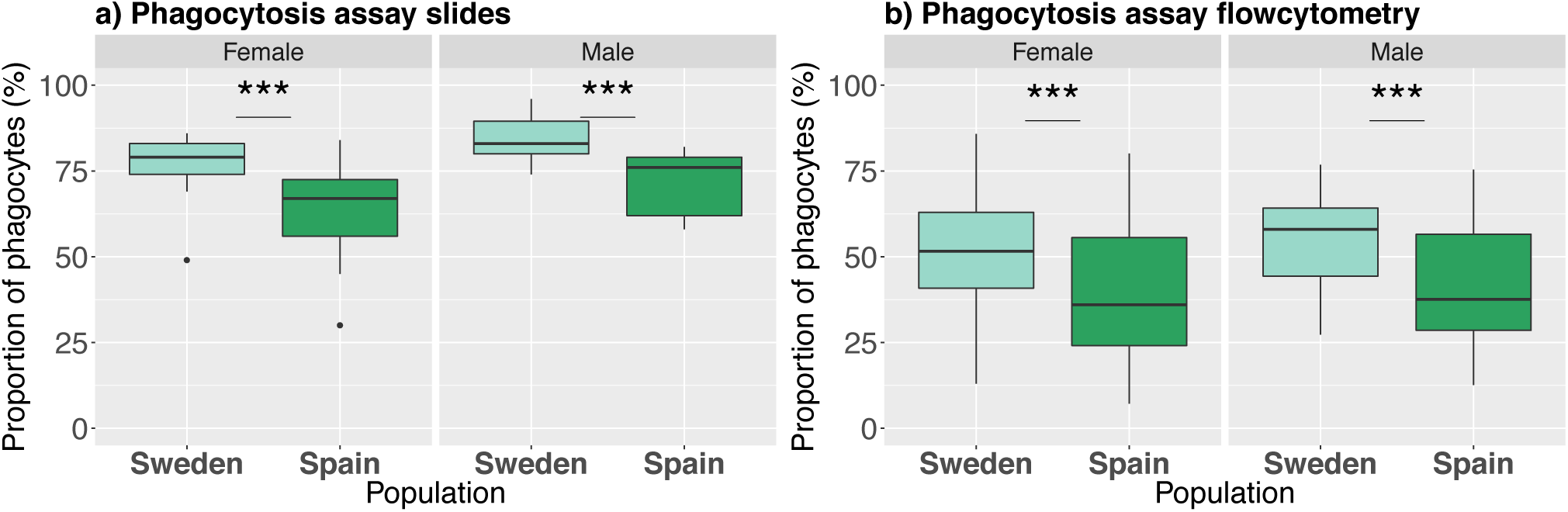
Proportion of phagocytes in *P. napi* larvae. (a) after injection of heat killed *E. coli*, calculated via imaging (N=44), and (b) after exposure to pHrodo conjugated E. coli, detected by flow cytometry (N=122). Bar plots are as described previously. Stars denote significance, note that for the phagocytosis slides males and females also significantly differed.

### Hemocyte composition and propensity of phagocytes

Phagocytic activity was detected in the three main cell types: granulocytes, plasmatocytes and early oenocytoids. The cell composition of these phagocytes did not significantly differ between the populations (proportion of phagocytic granulocytes: P = 0.52, SM Table 9; plasmatocytes: P = 0.60, SM Table 10; oenocytoids: P = 0.06, SM Table 11), although Swedish larvae appeared to have a trend towards a higher proportion of phagocytic oenocytoids compared to Spanish larvae (P= 0.06; Sweden M= 9.70% SD= 4.91 Spain M=7.33% SD= 3.27), similar to our previous observation of moderate differences in overall oenocytoid levels between populations (Fig 3c). However, the cell composition of the phagocytes differed significantly between the sexes, with females having a higher proportion of phagocytic granulocytes (Figure 5a; P = 0.013, SM table 9) and significantly fewer phagocytic plasmatocytes (Figure 5B; P = 0.0349, SM table 10), but not oenocytoids (Figure 5C; P = 0.086, SM Table 11).

**Figure 5.**
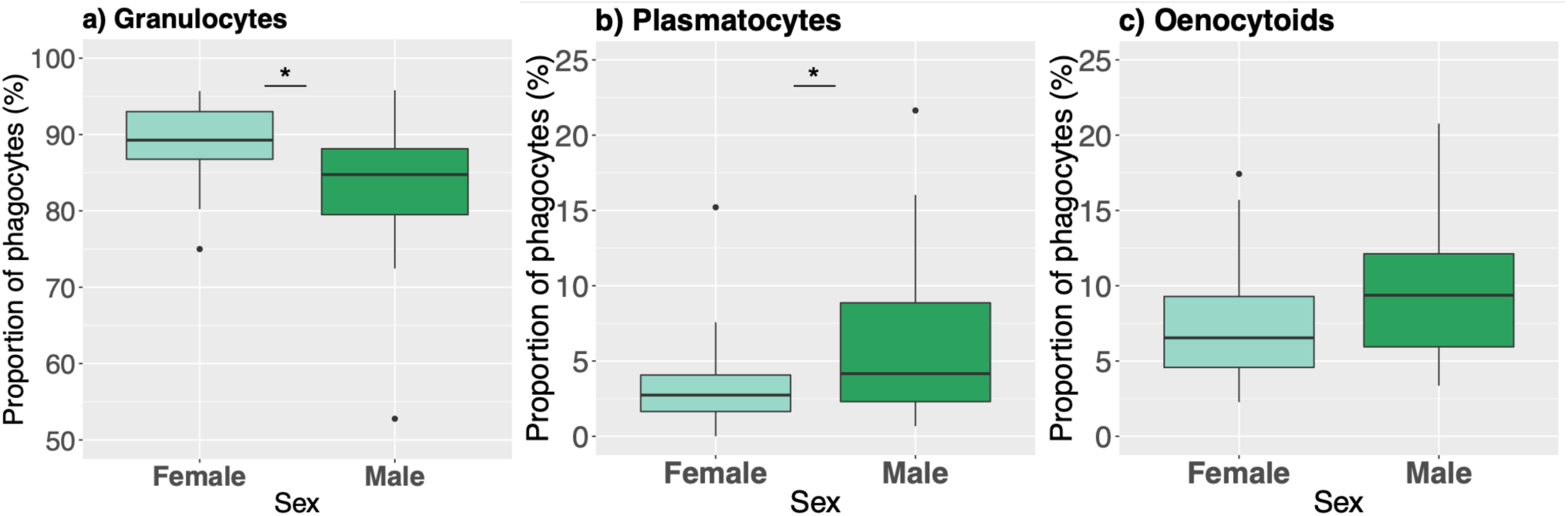
Differences between the sexes in cell composition of the phagocytes as measured via imaging (N=44). The majority of the phagocytes were granulocytes. Males had significantly more phagocytic plasmatocytes compared to females. Bar plots are as described previously.

Next, we investigated the propensity of a given cell type that was phagocytosing (i.e. the proportion of a given cell type to phagocytose). All three cell types showed a significantly higher propensity to phagocytose in Swedish larvae compared to Spanish larvae (GLM: P_granulocytes_ =0.00347, SM Table 12; P_plasmatocytes_ = 0.0018, SM Table 13; P_oenocytoids_ < 0.001, SM Table 14). In addition, males exhibited a higher propensity to phagocytose in all three cell types (GLM: P_granulocytes_ = 0.02442, SM Table 12; P_plasmatocytes_ = 0.0192, SM Table 13; P_oenocytoids_ = 0.002887, SM Table 14).

### Population differentiation

To explore the potential genetic basis for these differences in phagocytic capability, we investigated the divergence between the two populations in the 73 phagocytosis genes identified compared to the rest of the genome. First, F_ST_ values were calculated for each SNP within the coding region of genes. Genome-wide, a total of 363,459 exonic SNPs were identified with an average F_ST_ of 0.049 (SM table 15), and the genes involved with phagocytosis did not differ significantly from this genome wide pattern (Figure S1). Next, F_ST_ was estimated per gene (N= 13,692) to investigate the overall divergence patterns between the populations. Again, the genome-wide divergence between the population was relatively low (mean F_ST_ = 0.049; SM Table 15), and permutation tests revealed that the 73 phagocytosis genes were significantly less diverged compared to the other genes in the genome (mean F_ST_ = 0.038; SM Table 15, SM Figure 1). Finally, to investigate potential divergence in nearby regulatory regions, a second analysis investigated the 5kb up and downstream from each gene, with F_ST_ calculated at both the level of individual SNPs within this flanking region, as well as a single estimate for the entire region. Again, highly diverged regions of either analysis did not include phagocytosis genes (Figure 7). Finally, the top 2.5% outliers were selected for gene-set enrichment analysis (GSEA), which revealed that – somewhat in line with differences in phagocytic activities - F_ST_ outliers are enriched for genes involved with proteolysis, metabolic and catabolic processes, while no enrichment was found for phagocytosis per se (SM Tables 16, 17). Of note, while analyzing flanking regions, enrichment was repeatedly detected for genes involved in glutamine metabolism (SM Table 17).

**Figure 6.**
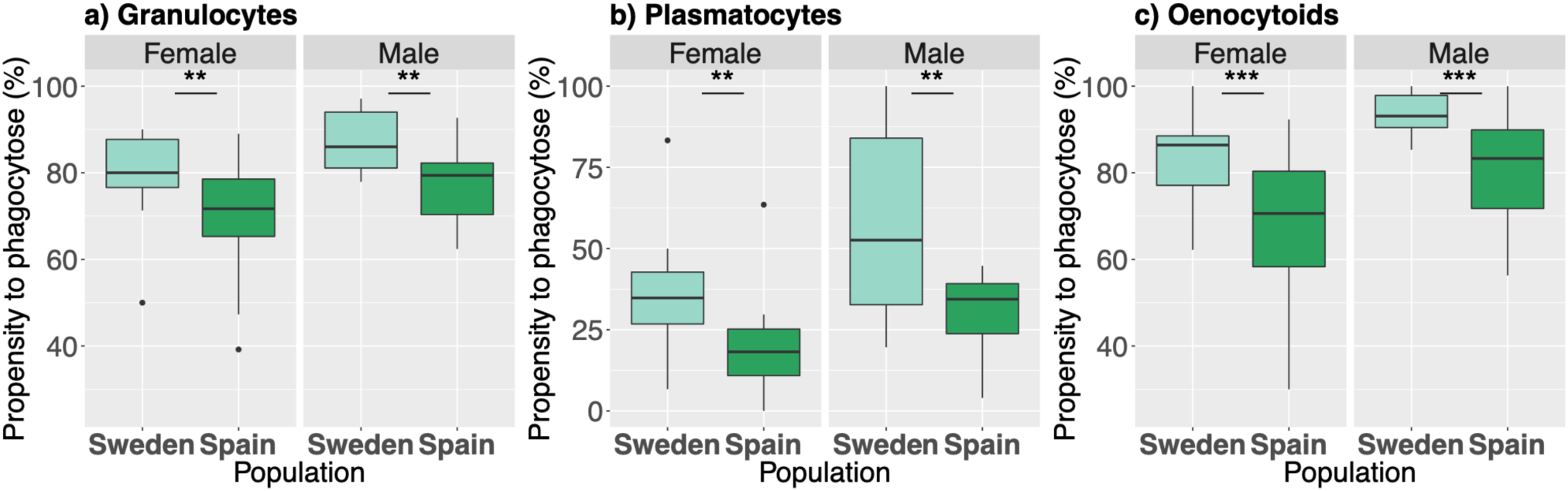
Propensity of a certain hemocyte type to phagocytose as measured via imaging (N=44). Populations differed in their cells propensity to phagocytose. In addition, the sexes also differed (p < 0.05*). Bar plots are as described previously. Stars denote significance. Note the differences in y-axis.

**Figure 7.**
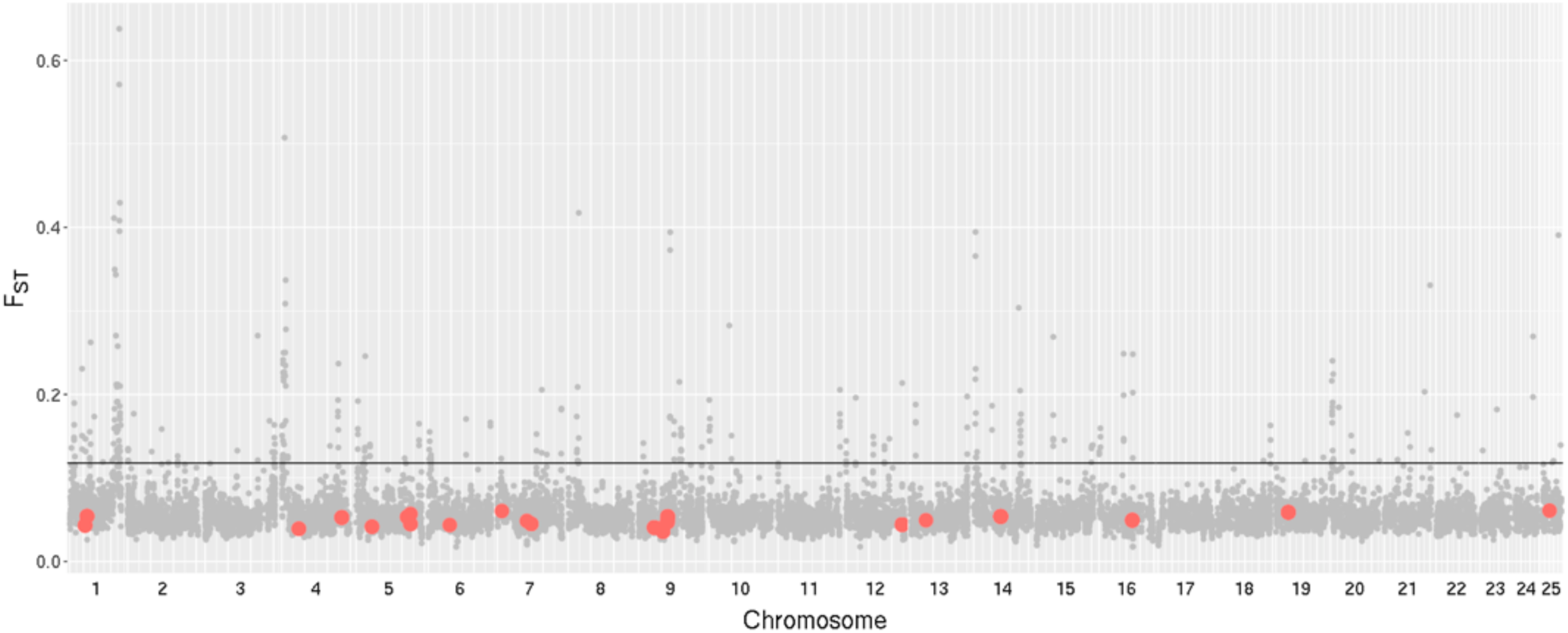
The genome-wide F_ST_ values of 5kb flanking regions, estimated per gene (n=13,692). Phagocytosis genes (n=73) are indicated (red dots), with many in close proximity. The grey line shows the 97.5 percentile of the analysis.

## Discussion

To date, few studies have explored the potential geographic variability in cellular immunity within species and between populations. Determining whether phagocytic capacity differs due to overall amount of cell types, their relative composition, or the phagocytic propensity of different cell types is challenging. Here we have investigated the hemocyte amount and composition, as well as phagocytic capacity in two allopatric populations of the green veined butterfly. We found that while the populations differed little in hemocyte amount and composition, they did differ significantly in their phagocytic capacity, which appears to be driven by increased level of their cells propensity to phagocytose. However, genome wide analysis of divergence between these populations found no excess genetic differentiation in genes annotated to phagocytic capacity, suggesting that our observed population differences might arise from genes affecting the activation and/or transdifferentiation of cells, which currently lack sufficient functional annotation.

### Species level patterns: Sex differences

According to Bateman’s principle, females are expected to invest more in post-copulation survival, since their fitness tends to be limited by their longevity, whereas males invest more in mating success. As a result, females are expected to invest more in their immune system (Rolff, 2002). Our results seem to be partly in concordance with this theory, although the total hemocyte count (THC) of unchallenged individuals did not differ between the sexes, the THC in females was significantly higher when inoculated with bacteria. This discrepancy could be a result of a sex difference in response to bacterial detection, wherein the females proliferate more hemocytes after infections compared to the males. However, in contrast, our data also reveal the opposite direction; males have more phagocytes (cells that engulfed bacteria) than females. This is consistent with previous studies in *P. napi*, wherein sex differences were observed in both directions: females have a higher encapsulation rate, whereas males exhibit higher phenoloxidase activity (Prasai and Karlsson 2011). Together, these results paint a complex picture, as it appears that there are differences between the sexes in their immune performance, but its direction depends on the immune phenotype studied. We did not measure whether the cell proliferation observed in females points towards an unquantified immune response, e.g. wound healing or phenoloxidase production, and it remains an open question.

### Population level variation: hemocyte composition, phagocytic capacity, and propensity

The populations differed in their cellular immunity in several ways. First, although the THC was not significantly different between the populations, the composition of the hemocytes did differ significantly. Spanish larvae contained a higher proportion of plasmatocytes, while Swedish larvae showed a trend for higher oenocytoid levels. Granulocytes, and to a lesser extent plasmatocytes, are reported to be the hemocytes responsible for phagocytosis in Lepidoptera (Ribeiro et al. 1996; Tojo et al., 2000; Lavine & Strand, 2002; Ling and Yu, 2006). However, our results reveal that early oenocytoids phagocytose at similar propensities as plasmatocytes. To our knowledge, this is a novel finding for Lepidoptera, though oenocytoids have been implicated in phagocytosis in several Coleopteran species (Giulianini et al., 2003; Giglio et al. 2008). Oenocytoids are the functional equivalent of *Drosophila* crystal cells and the early oenocytoids we observe may originate similar to novel crystal cells in *Drosophila* larvae, through transdifferentiation from plasmatocytes and/or granulocytes (Ribeiro & Brehelin, 2006; Leitao & Sucena, 2015; Anderl *et al*. 2016; Schmid et al. 2019). Second, we find a significant difference between the two populations in their overall phagocytic capability, as well as the propensity of individual cellular types to phagocytose. This could be a result of local adaptation to geographic variation in parasite type or load. Geographic variation in parasite pressure for butterflies has been observed before, as among adult monarchs captured at different points along the east coast fall migratory flyway, parasite prevalence declined as monarchs progressed southward (Bartel et al. 2011). Although geographic variation in parasite pressure is common among insect species, no data is available of the pathogenic environments of these two populations, and further research is needed to map this potential diversity to identify those of relevance to local adaption. Local adaptation implies a genetic basis for the difference between the populations. To assess this, we investigated the divergence between these populations in the genes directly annotated to be involved in phagocytosis.

### Population genomic analysis

Selection for differences in the immune performance between populations could act at several levels: the gene products (protein types), amount of products (protein levels), the cells that produce these proteins (cell proliferation), or the activation of cellular activity (cellular propensity). Despite observing a difference in phagocytosis, the genes previously characterized to be directly involved with phagocytosis (protein types) revealed no strong signal of genetic differentiation compared to the rest of the genome. Rather than selection acting directly upon the genes involved in phagocytosis itself, an alternative is that the phenotypic variation observed between populations has arisen through differences affecting the regulation, i.e. protein levels, of genes involved in phagocytosis. However, we find no excess divergence in the 5 kb flanking regions of phagocytosis genes. While differences in cell proliferation could give rise to our observations, our measures of THC and cell composition suggest this is not the case. Rather, our results may indicate that the selection has targeted either genes affecting the activation of phagocytic activity for each cell type, or a gene responsible for transdifferentiating cells, for example, plasmatocytes to oenocytoids, perhaps via a modulator gene. Unfortunately, given the lack of gene annotations for this function, this hypothesis remains untested at this time. A further alternative is that the genomic architecture is polygenic in nature rather than oligogenic, which would be difficult to detect given the diffuse nature of allelic changes that would then underlie our phenotypic changes between populations. It should be noted though that GSEA of flanking regions revealed differences in genes involved in glutamine metabolism (SM Table 17). Along with the Warburg effect, glutamine metabolic activities are increasingly implicated in activation of cells of both innate and adaptive immunity in mammals, including the differentiation of macrophages into M1 and M2 subtypes (Liu et al. 2017; Allison et al. 2017). One may speculate that similar metabolic changes contribute to transdifferentiation between hemocyte classes in insects although (Dolezal et al. 2019) – and not mutually exclusive – they are likely also crucial for additional physiological adjustments.

### Pleiotropic effects and phagocytosis

The interaction between immunity and other aspects of physiology suggests that natural selection on a trait might exert indirect pressure on other correlated fitness traits. Resistance to infection is often considered to be a result of the immune system, however, resistance involves the entire physiology of the host (Lazarro & Little, 2008). *P. napi* has an adaptive cline across latitude in several traits, one of which is the number of generations they produce in a year. The longer and warmer growing season in Spain permits up to four generations per year, whereas the shorter growing season in northern Sweden only allows one generation, and individuals spend the majority of their lifespan as diapausing (i.e. overwintering) pupae (Posledovich et al. 2015; Pruisscher et al., 2018). Overwintering insects face pathogen and parasite pressures that change with the seasons, and as a result are expected to invest more in their overall immunity (Ferguson and Sinclair, 2017). Secondly, immunity, longevity, reproduction, and metabolism are linked in a complex network via shared hormonal regulation (Flatt *et al*. 2005) and insects that have a relatively long life, are expected to invest in a more long-lasting sturdier body with more effective immune system (Karlsson and Wickman 1989; Prasai & Karlsson, 2011). Therefore, several indirect co-varying traits could be the basis of the variation in phagocytic capability found (i.e. the phagocytic propensity differences may arise indirectly via pleiotropy).

### Hemocyte and phagocytosis insights from multiple angles

Here we have measured hemocyte diversity and phagocytic capacity using several different approaches. Sometimes these have agreed, other times they have not. The sex-linked difference in THC was found using cell slides, but not detected by flow cytometry. This could be either due to the sample size used during the flow cytometer, or perhaps the type of cell that drives this difference is destroyed during this method. In sum, both methods have their strengths and potential biases. However, both methods are concordant in finding differences between populations in phagocytic capacity, while the sex level differences were unique to the slide assay, consistent with the latter being perhaps a smaller effect. A final consideration are the candidate genes used in our study, as these were identified via homology with *Drosophila* to be involved with phagocytosis. *P. napi l*acks experimental studies confirming their direct role of these genes, which is unfortunately a limitation common in non-model organisms.

## Conclusion

In sum, variation in the immune performance between populations can arise in various ways. We find evidence of differences in the phagocytic capability of populations, as well as the composition of the hemolymph. Cell types previously described to not be involved with phagocytosis, appear to have evolved the ability in our species. Our results suggest that to investigate genetic basis of phenotypic differences in immunity, candidate gene approaches can be limited in their insights, calling for the need for genome-wide, unbiased studies, perhaps using QTL or GWAS approaches.

## Acknowledgements

The authors would like to thank the various lab assistants for their help during the experiments. Ramprasad Neethiraj for general bioinformatic help, Sara Kurland for the statistics discussions, and Peter Pruisscher for helpful comments during writing. Funding was provided by the Swedish Research Council (2017-04386 and 621-2012-4001 to CWW, 2012-3715 and 2010-5341 to SN), and the Knut and Alice Wallenberg Foundation (grant number 2012.0058).

---

**SM Table 1.** Test Statistics for total hemocyte count by sex and population

|  | Estimate | Std. Error | t value | Pr(>\|t\|) |
| --- | --- | --- | --- | --- |
| (Intercept) | 15716 | 1927 | 8.156 | 1.31e-09 |
| PopulationSweden | -1853 | 2340 | -0.792 | 0.434 |
| SexMale | -1790 | 2350 | -0.762 | 0.451 |

**SM Table 2.** Test Statistics for total number of Phagocytosing cells

| Fixed effects: Tot.no.of.cells ~ Population + Sex |  |  |  |  |  |
| --- | --- | --- | --- | --- | --- |
|  | Value | Std.Error | DF | t-value | p-value |
| (Intercept) | 48.95381 | 6.345961 | 306 | 7.714168 | 0.0000 |
| PopulationSpain | 12.63784 | 7.328444 | 41 | 1.724492 | 0.0922 |
| SexM | -19.04693 | 7.328444 | 41 | -2.599042 | 0.0129 |

**SM Table 3.** Averages of proportion of phagocytic cell types by sex and population in percentages M= average, SD = standard deviation

|  | Sweden |  |  |  | Spain |  |  |  |
| --- | --- | --- | --- | --- | --- | --- | --- | --- |
|  | Female |  | Male |  | Female |  | Male |  |
|  | M | SD | M | SD | M | SD | M | SD |
| **Phagocytic granulocytes (%)** | 89.05 | 6.51 | 81.83 | 5.91 | 88.92 | 3.61 | 84.76 | 11.97 |
| **Phagocytic plasmacytes (%)** | 3.21 | 2.15 | 6.52 | 6.32 | 3.82 | 4.29 | 7.83 | 8.48 |
| **Phagocytic oenocytoids (%)** | 7.75 | 5.07 | 11.66 | 4.05 | 7.27 | 2.48 | 7.39 | 4.04 |

**SM Table 4.** Test Statistics for proportion of Granulocytes between populations and sex

| Fixed effects: Prop_Gran ~ Population + Sex |  |  |  |  |  |
| --- | --- | --- | --- | --- | --- |
|  | Value | Std.Error | DF | t-value | p-value |
| (Intercept) | 0.8500781 | 0.02208492 | 306 | 38.49134 | 0.0000 |
| PopulationSpain | -0.0325426 | 0.02550832 | 41 | -1.27576 | 0.2092 |
| SexM | -0.0488211 | 0.02550832 | 41 | -1.91393 | 0.0626 |

**SM Table 5.**
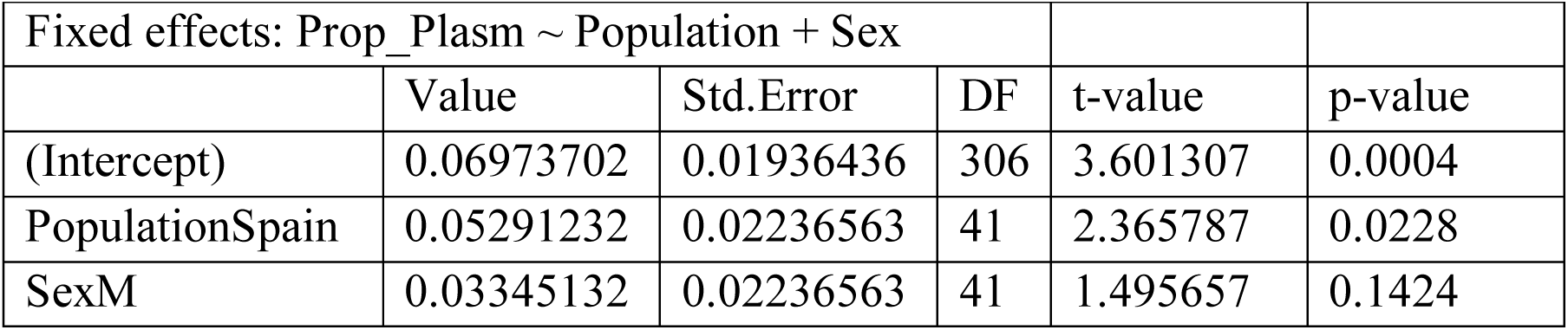
Test Statistics for proportion of Plasmatocytes between populations and sex

**SM Table 6.** Test Statistics for proportion of Oenocytoids between populations and sex

| Fixed effects: Prop_Oeno ~ Population * Sex |  |  |  |  |  |
| --- | --- | --- | --- | --- | --- |
|  | Value | Std.Error | DF | t-value | p-value |
| (Intercept) | 0.06977273 | 0.01074684 | 306 | 6.492396 | 0.0000 |
| PopulationSpain | 0.00102273 | 0.01519832 | 40 | 0.067292 | 0.9467 |
| SexM | 0.03746586 | 0.01522313 | 40 | 2.461113 | 0.0183 |
| PopulationSpain:SexM | -0.04371586 | 0.02151123 | 40 | -2.032235 | 0.0488 |

**SM Table 7.**
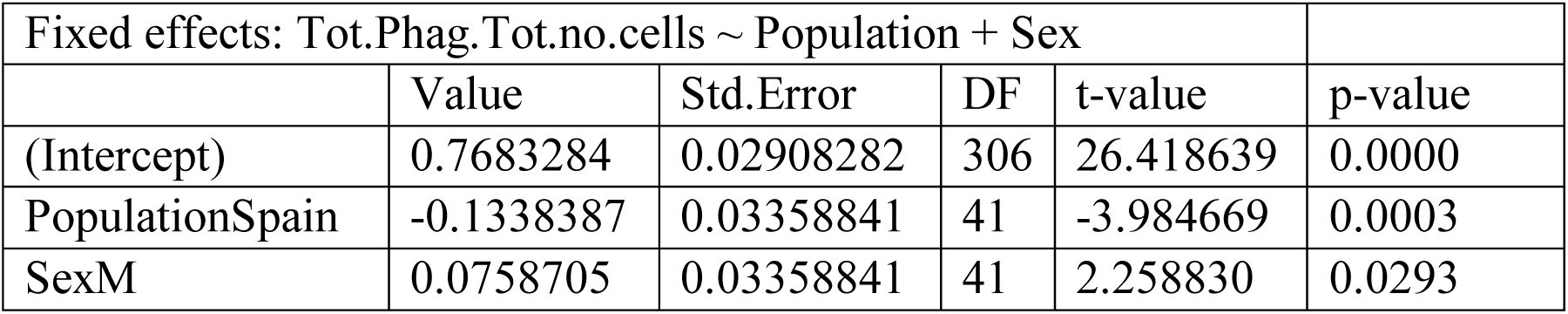
Test Statistics for total number of phagocytosing cells between population and sex

**SM Table 8.** Test statistics for proportion of phagocytes between population and sex as measured by flowcytometry

|  | Estimate | Std.Error | tvalue | Pr(>\|t\|) |
| --- | --- | --- | --- | --- |
| (Intercept) | -0.28436 | 0.10548 | -2.696 | 0.008040 |
| PopulationSweden | 0.49947 | 0.13242 | 3.772 | 0.000254 |
| SexMale | 0.01866 | 0.12947 | 0.144 | 0.885617 |

**SM Table 9.**
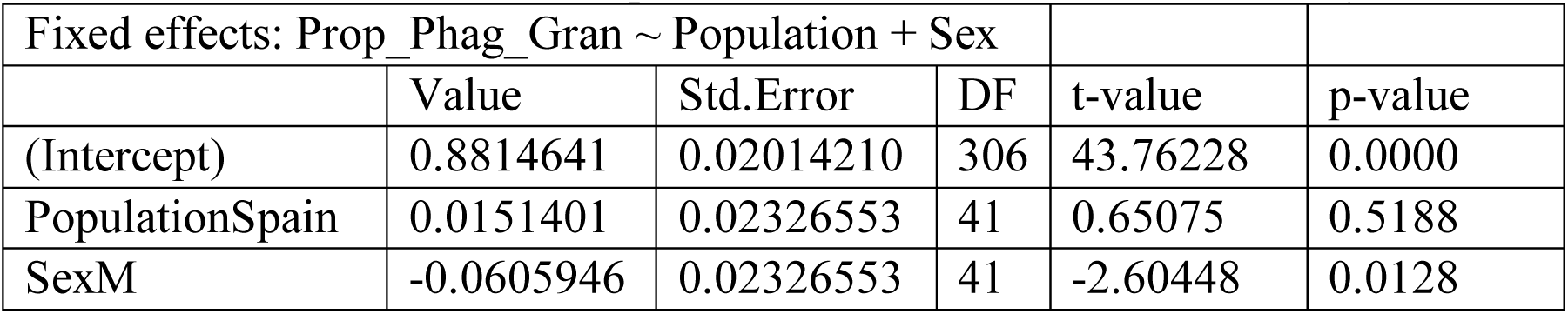
Test Statistics for proportion of phagocytosing granulocytes

| Fixed effects: Prop_Phag_Gran ~ Population + Sex |  |  |  |  |  |
| --- | --- | --- | --- | --- | --- |
|  | Value | Std.Error | DF | t-value | p-value |
| (Intercept) | 0.8814641 | 0.02014210 | 306 | 43.76228 | 0.0000 |
| PopulationSpain | 0.0151401 | 0.02326553 | 41 | 0.65075 | 0.5188 |
| SexM | -0.0605946 | 0.02326553 | 41 | -2.60448 | 0.0128 |

**SM Table 10.** Test Statistics for proportion of phagocytosing plasmatocytes

| Fixed effects: Prop_Phag_Plasm ~ Population + Sex |  |  |  |  |  |
| --- | --- | --- | --- | --- | --- |
|  | Value | Std.Error | DF | t-value | p-value |
| (Intercept) | 0.03105434 | 0.01494720 | 306 | 2.0776024 | 0.0386 |
| PopulationSpain | 0.00914131 | 0.01726409 | 41 | 0.5294985 | 0.5993 |
| SexM | 0.03767687 | 0.01726409 | 41 | 2.1823837 | 0.0349 |

**SM Table 11.** Test Statistics for proportion of phagocytosing oenocytoids

| Fixed effects: Prop_Phag_Oeno ~ Population + Sex |  |  |  |  |  |
| --- | --- | --- | --- | --- | --- |
|  | Value | Std.Error | DF | t-value | p-value |
| (Intercept) | 0.08854209 | 0.01096815 | 306 | 8.072658 | 0.0000 |
| PopulationSpain | -0.02447054 | 0.01267218 | 41 | -1.931044 | 0.0604 |
| SexM | 0.02231145 | 0.01267218 | 41 | 1.760663 | 0.0858 |

**SM Table 12.** Test Statistics for propensity of granulocytes to perform phagocytosis

| Estimate | Std. Error | t value | Pr(>\|t\|) |  |
| --- | --- | --- | --- | --- |
| (Intercept) | 1.3910 | 0.1704 | 8.162 | 3.98e-10 |
| PopulationSpain | -0.6058 | 0.1953 | -3.103 | 0.00347 |
| SexM | 0.4533 | 0.1940 | 2.337 | 0.02442 |

**SM Table 13.** Test Statistics for propensity of plasmatocytes to perform phagocytosis

| Estimate | Std. Error | t value | Pr(>\|t\|) |  |
| --- | --- | --- | --- | --- |
| (Intercept) | -0.4672 | 0.2461 | -1.899 | 0.0647 |
| PopulationSpain | -0.9893 | 0.2963 | -3.339 | 0.0018 |
| SexM | 0.7188 | 0.2949 | 2.437 | 0.0192 |

**SM Table 14.**
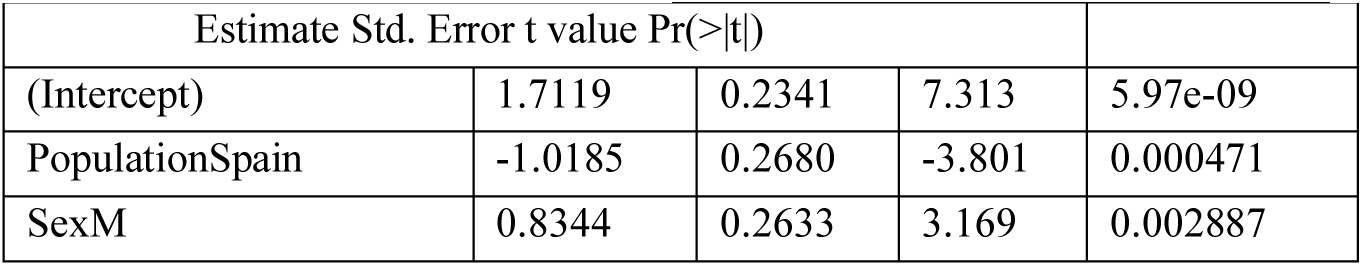
Test Statistics for propensity of oenocytoids to perform phagocytosis

| Estimate | Std. Error | t value | Pr(>\|t\|) |  |
| --- | --- | --- | --- | --- |
| (Intercept) | 1.7119 | 0.2341 | 7.313 | 5.97e-09 |
| PopulationSpain | -1.0185 | 0.2680 | -3.801 | 0.000471 |
| SexM | 0.8344 | 0.2633 | 3.169 | 0.002887 |

**SM Table 15.** Distribution of F_ST_ values G stands for genome-wise values, whereas P is the phagocytosis genes values

| Percentile | SNPs |  |  |  | Gene level |  |  |  |
| --- | --- | --- | --- | --- | --- | --- | --- | --- |
| | $F_{ST}$ within genes | | $F_{ST}$ 5kb genes | | $F_{ST}$ within genes | | $F_{ST}$ 5kb genes | |
|  | G | P | G | P | G | P | G | P |
| 100% | 1 | 0,308 | 1 | 0.714 | 0.795 | 0.142 | 0.638 | 0.079 |
| 99.5% | 0.329 | 0,256 | 0.350 | 0.273 | 0.268 | 0.142 | 0.201 | 0.079 |
| 97.5% | 0.191 | 0,191 | 0.204 | 0.178 | 0.141 | 0.097 | 0.118 | 0.064 |
| 75% | 0.065 | 0,068 | 0.065 | 0.063 | 0.059 | 0.060 | 0.063 | 0.054 |
| 50% | 0.034 | 0,032 | 0.033 | 0.033 | 0.043 | 0.041 | 0.053 | 0.049 |
| 2.5% | 0.002 | 0,003 | 0.002 | 0.003 | 0 | 0 | 0.033 | 0.036 |
| 0% | 0 | 0,001 | 0 | 0 | 0 | 0 | 0 | 0.036 |
| Average | 0.049 | 0,049 | 0.050 | 0.047 | 0.049 | 0,038 | 0.058 | 0.049 |
| Std Dev | 0.057 | 0,049 | 0.059 | 0.049 | 0.042 | 0,027 | 0.026 | 0.007 |

**SM Table 16.**
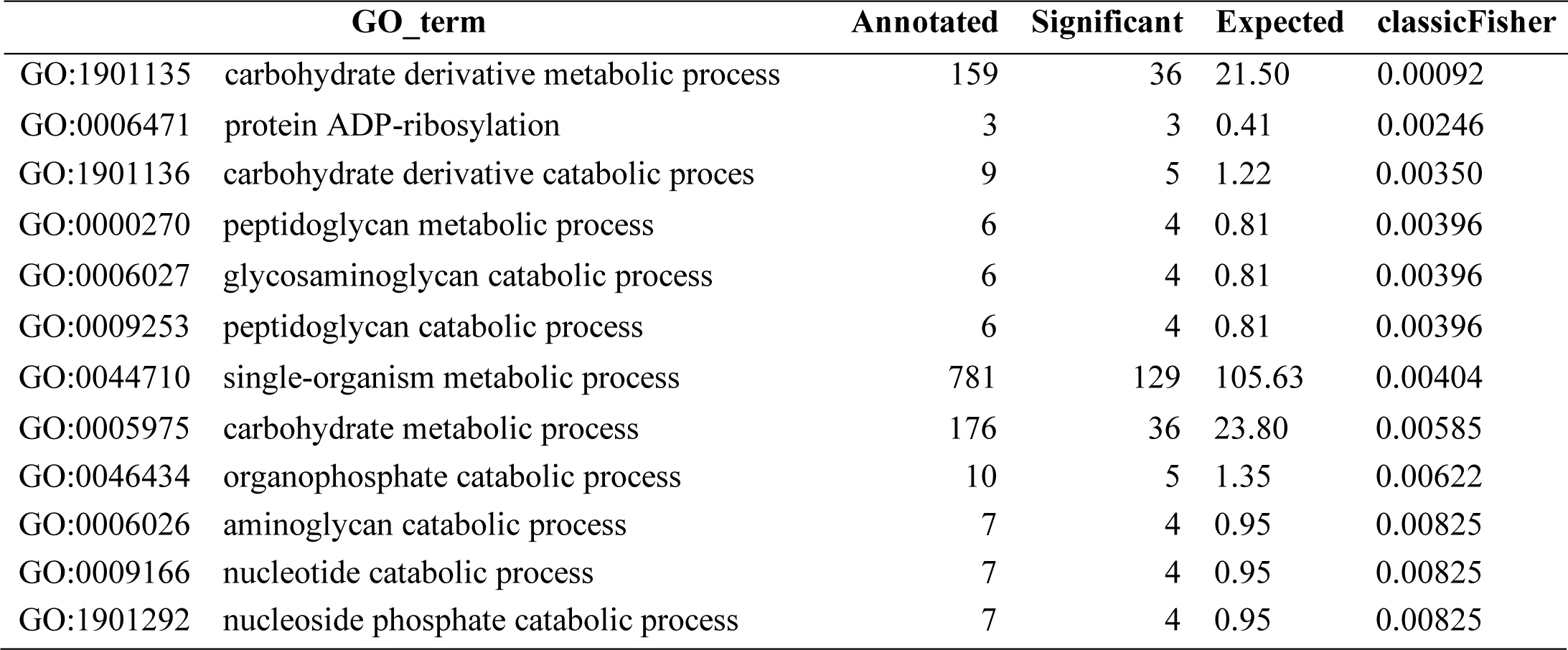
The significant GO terms with p < 0.01 of the SNPs with the highest 2.5 % values (exons only).

| GO_term | Annotated | Significant | Expected | classicFisher |
| --- | --- | --- | --- | --- |
| GO:1901135 carbohydrate derivative metabolic process | 159 | 36 | 21.50 | 0.00092 |
| GO:0006471 protein ADP-ribosylation | 3 | 3 | 0.41 | 0.00246 |

|  |  |  |  |  |  |
| --- | --- | --- | --- | --- | --- |
| GO:1901136 | carbohydrate derivative catabolic proces | 9 | 5 | 1.22 | 0.00350 |
| GO:0000270 | peptidoglycan metabolic process | 6 | 4 | 0.81 | 0.00396 |
| GO:0006027 | glycosaminoglycan catabolic process | 6 | 4 | 0.81 | 0.00396 |
| GO:0009253 | peptidoglycan catabolic process | 6 | 4 | 0.81 | 0.00396 |
| GO:0044710 | single-organism metabolic process | 781 | 129 | 105.63 | 0.00404 |
| GO:0005975 | carbohydrate metabolic process | 176 | 36 | 23.80 | 0.00585 |
| GO:0046434 | organophosphate catabolic process | 10 | 5 | 1.35 | 0.00622 |
| GO:0006026 | aminoglycan catabolic process | 7 | 4 | 0.95 | 0.00825 |
| GO:0009166 | nucleotide catabolic process | 7 | 4 | 0.95 | 0.00825 |
| GO:1901292 | nucleoside phosphate catabolic process | 7 | 4 | 0.95 | 0.00825 |

**SM Table 17.** The GO terms with terms with p < 0.01 of the SNPs with the highest 2.5 % F ST values (with 5kb both sides)

|  | GO_term | Annotated | Significant | Expected | classicFisher |
| --- | --- | --- | --- | --- | --- |
| GO:0006508 | proteolysis | 380 | 45 | 14.90 | 1.6e-12 |
| GO:0008152 | metabolic process | 2436 | 119 | 95.54 | 8.4e-06 |
| GO:0006542 | glutamine biosynthetic process | 5 | 3 | 0.20 | 0.00056 |
| GO:0044710 | single-organism metabolic process | 781 | 46 | 30.63 | 0.00154 |
| GO:0019538 | protein metabolic process | 872 | 50 | 34.20 | 0.00162 |
| GO:0006541 | glutamine metabolic process | 7 | 3 | 0.27 | 0.00184 |
| GO:0055114 | oxidation-reduction process | 391 | 26 | 15.34 | 0.00434 |
| GO:0008652 | cellular amino acid biosynthetic process | 19 | 4 | 0.75 | 0.00555 |
| GO:1901607 | alpha-amino acid biosynthetic process | 19 | 4 | 0.75 | 0.00555 |
| GO:0009084 | glutamine family amino acid biosynthetic... | 11 | 3 | 0.43 | 0.00773 |
| GO:0009064 | glutamine family amino acid metabolic pr... | 13 | 3 | 0.51 | 0.01265 |
| GO:0016053 | organic acid biosynthetic process | 28 | 4 | 1.10 | 0.02233 |
| GO:0046394 | carboxylic acid biosynthetic process | 28 | 4 | 1.10 | 0.02233 |
| GO:0006265 | DNA topological change | 7 | 2 | 0.27 | 0.02818 |
| GO:1901605 | alpha-amino acid metabolic process | 33 | 4 | 1.29 | 0.03841 |

**Figure S1.**
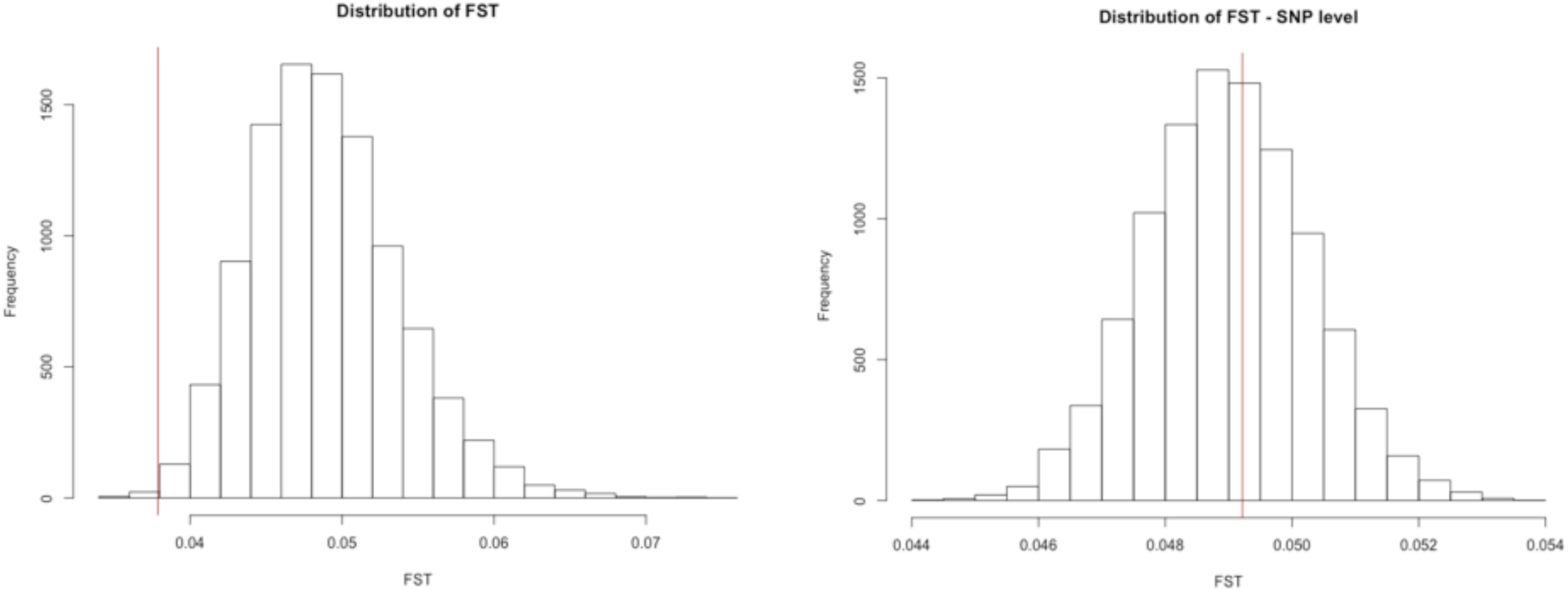
Distribution plots of the permutations for genome-wide FST, left calculated per gene, right per SNP. The red line denotes the averages of these values for the phagocytosis genes.

**Figure S2 - Supplementary figure 2.**
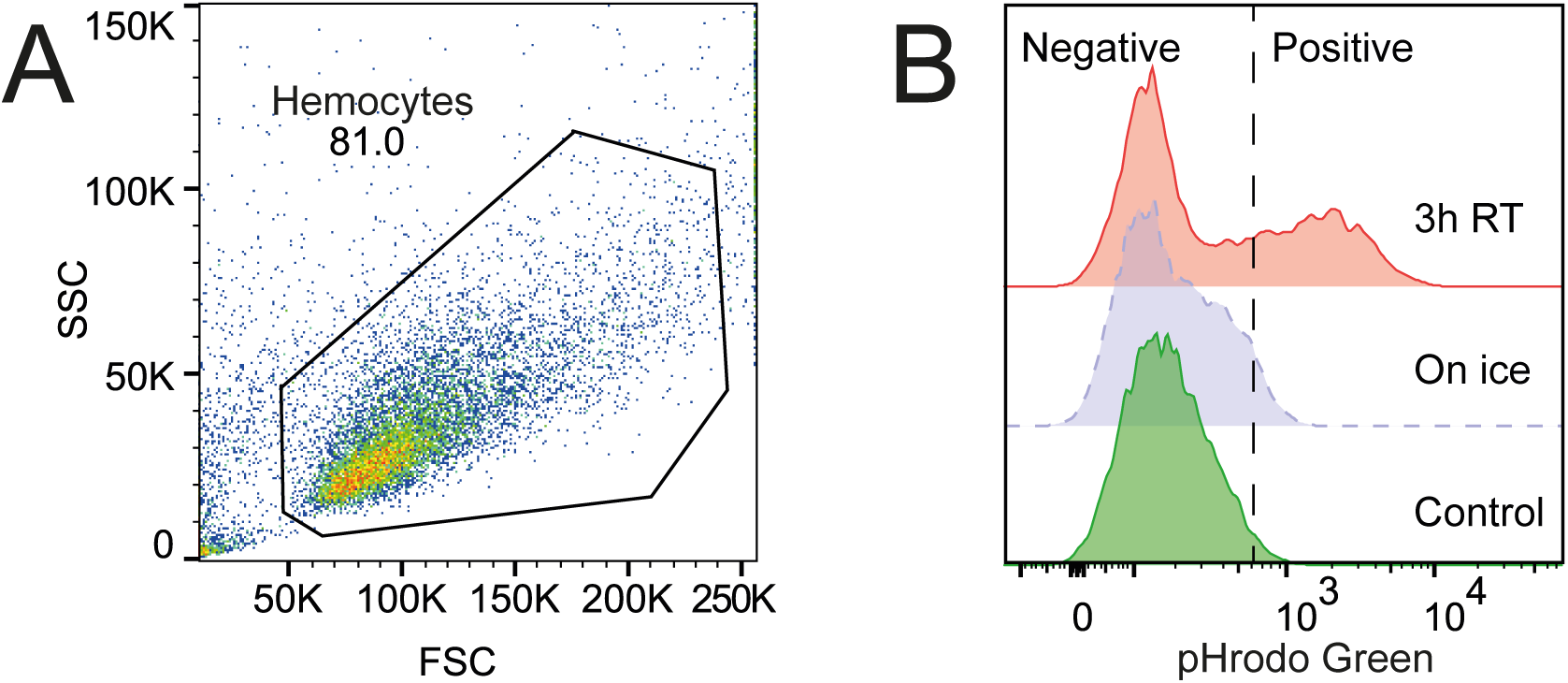
Gating strategy and example of staining for the flow cytometry assay (A) Hemocytes from *in vitro* samples were selected and gated on forward (FSC) and side scatter (SSC) to exclude debris. (B) Histograms showing examples of uptake of E.coli in hemocytes. Phagocytic activity of pHrodo™ Green E. coli was measured by fluorescence, and threshold was set according to samples incubated on ice (dashed line). Hemocytes that stained above this threshold were put as positive for phagocytic activity of E. coli.

